# Anti-cancer potential of cannabis terpenes in a taxol-resistant model of breast cancer

**DOI:** 10.1101/2021.10.08.463667

**Authors:** Andrea M. Tomko, Erin G. Whynot, Lauren F. O’Leary, Denis J. Dupré

**Affiliations:** Department of Pharmacology, Dalhousie University, Halifax, NS, Canada

**Keywords:** breast cancer, terpenes, chemotherapeutic resistance, cannabis, apoptosis, synergy

## Abstract

Chemotherapeutic resistance can limit breast cancer outcomes; therefore, the exploration of novel therapeutic options is warranted. Isolated compounds found in cannabis have previously been shown to exhibit anti-cancer effects, but little is known about their effects in resistant breast cancer. Our study aims to evaluate the effects of terpenes found in cannabis in *in vitro* chemotherapy-resistant model of breast cancer. We aimed to identify whether five terpenes found in cannabis produced anti-cancer effects, and if their effects were improved upon co-treatment with cannabinoids and flavonoids also found in cannabis. Nerolidol and β-caryophyllene produced the greatest cytotoxic effects, activated the apoptotic cascade and reduced cellular invasion. Combinations with the flavonoid kaempferol potentiated the cytotoxic effects of ocimene, terpinolene, and β-myrcene. Combinations of nerolidol and Δ9-tetrahydrocannabinol or cannabidiol produced variable responses ranging from antagonism and additivity to synergy, depending on concentrations used. Our results indicate that cannabis terpenes, alone or combined with cannabinoids and flavonoids, produced anti-cancer effects in chemotherapy-resistant breast cancer cell lines. This study is a first step in the identification of compounds that could have therapeutic potential in the treatment of resistant breast cancer.

## Introduction

Breast cancer has an incidence rate of 24% and a mortality rate of 15% in women, globally (Bray et al., 2018). Although survival has improved due to treatment options such as chemotherapy, radiation, and surgery, these treatments can be of limited use for various reasons, such as metastasis, which renders surgical removal non-viable (Morris et al., 2009). Another significant limitation to current anti-cancer therapy is chemotherapeutic resistance that limits the effectiveness of chemotherapy drugs and reduces survival (Longley & Johnston, 2005; Marquette & Nabell, 2012). There is a need for novel treatments that overcome or evade cellular resistance to chemotherapeutic agents to provide a wider range of therapeutic options and improve outcomes for patients.

The Cannabis plant is a polypharmacy that comprises hundreds of compounds including phytocannabinoids, terpenes, and flavonoids, and some of these compounds have been shown to exert anti-cancer effects (Klumpers & Thacker, 2019; Tomko et al., 2020). Terpenes are aromatic compounds that are present in lower amounts than cannabinoids, typically ranging between 1-5% of a particular cultivar of cannabis with profiles varying by cultivar and some associated with varying Δ9-tetrahydrocannbinol (THC) or cannabidiol (CBD) content (Chan et al., 2016; Klumpers & Thacker, 2019; Lewis et al., 2018). Terpenes’ role as insecticidal and antibacterial agents in plants suggest a range of effects that has led researchers to investigate their results in humans where they have shown anti-inflammatory, anti-parasitic, anti-bacterial, and anti-cancer properties (Chan et al., 2016; Tetali, 2019). Sesquiterpenes found in cannabis have shown anti-cancer effects in *in vitro* and *in vivo* models of cancer including induction of apoptosis, cell cycle arrest and improving sensitivity to several chemotherapeutic agents (Tomko et al., 2020). Along with the cannabinoids and terpenes, about 25 flavonoids have been identified from 7 chemical structures in cannabis (Andre et al., 2016; Elsohly & Slade, 2005; Radwan et al., 2008; Segelman et al., 1977). Flavonoids may impact one or several of the various mechanisms involved in the regulation of multidrug resistance such as efflux pump expression and function, oxidative stress, or signaling pathways involved in cellular viability, in addition to their ability to directly inhibit the proliferation of cancer cells (Tomko et al., 2020; Ye et al., 2019).

Although individual compounds found in cannabis exhibit anti-cancer properties (Ambrož et al., 2017; Hanušová et al., 2017), it has been suggested that whole plant extracts may have greater anti-cancer effects than individual components, potentially due to complementary or synergistic effects between the compounds found in the plant through the activation of various complementary signaling pathways (Blasco-Benito et al., 2018). While some information is available regarding the effects of terpenes and flavonoids on cancer cell lines, they remain largely uncharacterized in terms of their effects on chemotherapy-resistant tumours. We and others have demonstrated that some cannabinoids evade resistance mechanisms involved in resistance to taxol (Barbado et al., 2017; Holland et al., 2006, 2007; Tomko et al., 2019) and it is possible that other compounds present in Cannabis can act similarly. For example, several terpenes found in plants have biological actions including anti-cancer effects (Chan et al., 2016; Cho et al., 2017)(Chan et al., 2016), however their effects in multidrug resistant cells have yet to be discovered. Due to the potential role of flavonoids, terpenes and cannabinoids in various hallmarks of cancer, we investigated their anti-cancer potential when combined in multidrugresistant cancer cell lines. We showcase here the anti-cancer activity of several terpenes in paclitaxel(taxol)-resistant (PR) models of breast cancer, as well as their potential effects when combined with the flavonoid kaempferol and the cannabinoids THC and CBD. Our results suggest that Nerolidol (Ner) and β-caryophyllene (BC) inhibit the proliferation and invasion of PR breast cancer cells and show potential to enhance effects when combined with kaempferol, THC, or CBD.

## Methods

### Compounds

Paclitaxel, Δ9-tetrahydrocannbinol (THC) and cannabidiol (CBD) were purchased from Sigma-Aldrich. Nerolidol (Ner), β-caryophyllene (BC), Ocimene (Oci), β-myrcene (BM), Terpinolene (Terp) and Kaempferol were purchased from Cayman Chemical Company.

### Cell Culture

Human Breast Adenocarcinoma MDA-MB-231 and MCF-7 cells were cultured in Dulbecco’s Modified Eagle’s Medium-high glucose (DMEM) with 1% penicillin-streptomycin containing 10% fetal bovine serum (FBS). PR MDA-MB-23 cells (470nM) were obtained from Dr. Kerry Goralski, Dr. David Hoskin and Anna Greenshields, Dalhousie University. PR MCF-7 cells (470nM) were obtained from Dr. Robert Robey and Susan Bates, National Cancer Institute. PR MDA-MB-231 cells and PR MCF-7 cells were cultured in DMEM with 1% pen-strep, 10% FBS and 470nM Paclitaxel. Non-tumorigenic breast epithelial MCF10-A cells were cultured in DMEM/F12 media containing 5% horse serum, 0.5mg/ml hydrocortisone, 100ng/ml cholera toxin, 10μg/ml insulin and 1% penicillin-streptomycin. All cells were incubated at 37°C in 5% CO_2_.

### Cytotoxicity Assay

MDA-MB-231, MCF-7 cells and their PR equivalents and MCF10-A cells were seeded at 10,000 cells per well in black 96 well plates and incubated at 37° at 5% CO_2_ for 24h. Cells were treated with concentrations of Ner, BC, Oci, BM, or Terp alone or in combination with kaempferol using DMSO as a vehicle control, with 1% FBS and incubated for 24h. Drug treatments were repeated after 24h for a total time of 48h. AlamarBlue was added to each well equal to 10% of the total volume per well and incubated for 3h. Fluorescence was measured at 560nm excitation and 590nm emission using a Biotek Cytation 3 plate reader. Cell viability was calculated as a percent relative to the vehicle control and presented as mean ± SEM.

### Annexin V Apoptosis Assay

MDA-MB-231 and MCF-7 cells were seeded at 5,000 cells/well in a 96 well plate and incubated for 24h. Cells were treated with 5μM Ner, BC, or DMSO with 1% FBS and incubated for 24h. Cells were resuspended in annexin V assay buffer and incubated in the dark with propidium iodide (PI) and annexin V–fluorescein isothiocyanate–conjugated stain for 20 min. Cells were observed by fluorescence microscopy, and a minimum of five fields of view were recorded using an Olympus IX81 microscope with a Photometrics coolSNAP HQ2 camera and an Excite series 120Q light source. Annexin V stain was excited at 488 nm and imaged at 525 nm and propidium iodide was excited at 535 nm and imaged at 617 nm. Rates of early apoptosis were determined by dividing the number of cells that stained positive for annexin V divided by the total number of cells (Young et al., 2015; Martin et al., 2019).

### Caspase 3/7 Apoptosis Assay

MDA-MB-231 and MCF-7 cells were seeded in black 96 well plates at 10,000 cells/ well and incubated for 24h. Cells were treated with 5μM Ner, BC, or DMSO with 1% FBS and incubated for 24h. Media was removed from wells and 100 μl of 5μM Cell Event Caspase 3/7 Green Detection Reagent in 1X phosphate-buffered saline (PBS) with 5% FBS was added to each well. Following a 30-minute incubation fluorescence was read at 488 nm excitation and 530 nm emission using a using a Biotek Cytation 3 plate reader. Cells with activated caspase 3/7 were quantified as a percent relative to the vehicle control and presented as mean ± SEM.

### Cell Lysis and Western Blotting

MDA-MB-231 cells were seeded at 100,000 cells/well in 6 well plates and incubated for 24h. Cells were treated with 5μM Ner, BC, or DMSO with 1% FBS for 24h. Cells were detached, pelleted by centrifugation, and lysed using 150 μl RIPA buffer (150 mM NaCl, 50 mM Tris-HCl pH 7.5, 1% NP40, 0.5% sodium deoxycholate, 0.1% sodium dodecyl sulfate, and Roche’s cOmplete™ EDTA-free protease inhibitor cocktail. Bovine serum albumin–coated Protein A-Sepharose beads and 10% DNase I were added to remove nucleic acid and organellar material from the sample. Lysates were mixed 50:50 with Laemmli buffer containing 5% 2-mercaptoethanol. Samples were run on a sodium dodecyl sulfate–polyacrylamide electrophoresis gel, transferred to a nitrocellulose membrane and blocked in a PBS solution containing 10% skim milk powder for 60 minutes. Primary antibodies (beta-Actin and HRP-conjugated Caspase-3 (Santa Cruz Biotechnology) were added to the milk solution (1:1000) and incubated overnight at 4°. Membranes were washed with TBS-Tween and incubated in secondary antibodies for 1h (1:1000). Membranes were washed again with TBS-Tween to remove any unbound antibody. Chemiluminescence was performed using Western Lightning® Plus-ECL Enhanced Chemiluminescence Substrate (PerkinElmer) and then membranes were exposed to x-ray film and developed for 1-10 minutes.

### Matrigel Invasion Assay

Growth factor reduced 8.0-μm Matrigel Invasion Chambers (Corning), and Cell Culture Inserts with an 8.0-μm membrane were added to a 24-well plate. Matrigel Invasion Chambers were hydrated with 250 μL of DMEM containing 0.2% FBS, and 10 μM of Ner, BC or DMSO and incubated for 1h at 37°C. Following incubation, 700 μL of DMEM containing 10% FBS was added to the lower chamber of Matrigel Invasion Chambers and Cell Culture Inserts and 250 μl of DMEM containing 0.2% FBS DMSO was added to the upper chamber of cell culture inserts. Two hundred and fifty microliters of MDA-MB-231 cells at 100,000 cells/mL was added to each cell culture insert and Matrigel invasion chamber resulting in a final cell concentration of 25,000 cells/well and drug concentration of 5 μM. Cells were incubated for 24h. After incubation, cells that did not invade through matrigel or migrate were removed from the inside of the insert using a cotton swab dampened with PBS. Wells were fixed in methanol for 10 min and then stained with 3.5 g/L crystal violet in 2% ethanol solution for 10 minutes. Following staining, wells were rinsed with H_2_O and left to dry overnight. Cells that migrated or invaded through the membranes were counted using an Olympus CKX41 light microscope. The number of cells invaded for each condition were represented as a percentage relative to the number of cells invaded when exposed to the vehicle control.

### Verapamil Efflux Pump Assay

PR MDA-MB-231 and PR MCF-7 cells were seeded at 10,000 cells/well and incubated at 37° C and 5% CO_2_ for 24h. Cells were treated with verapamil alone, DMSO, or 0.1μM of each of the terpenes +/- 10 μM verapamil, with 1% FBS. Cells were incubated for 24h, and drug treatments were repeated, for a total of 48h. Fluorescence was quantified as previously described using AlamarBlue® after 48 h treatment.

### Assessment of Synergy, Additivity, and Antagonism

Synergy between THC or CBD and Ner were evaluated using the checkerboard assay in PR MDA-MB-231 cells. Cells were seeded 10,000 cells per well in black 96-well plates with a final volume of 100 μL per well. Cannabinoid concentrations ranged from 0-10μM and Ner concentrations from 0-25μM. Fluorescence was quantified as previously described using AlamarBlue® after 48 h treatment. The analysis was performed using SynergyFinder 2.0 (Ianevski et al., 2020), where the Bliss independence drug interaction model was used. Drug combination responses were also plotted as 3D synergy maps using SigmaPlot to assess the potential synergy, antagonism or additive behaviours of the drug combinations. The summary synergy represents the average excess response due to drug interactions. A synergy score of <-10 was considered as antagonistic, a range from −10 to +10 as additive and >+10 as synergistic.

### Statistical Analysis

Statistical analysis was completed using GraphPad Prism software. All error bars are representative of mean ± SEM. Unpaired student’s t-tests were performed for analysis of two independent groups. One-way ANOVA with Tukey’s post-hoc test was used to assess multigroup comparisons. p values are reported as follows: * p < 0.05, ** p < 0.01, *** p < 0.001.

## Results

### Cytotoxicity

We first examined the effects of BC, Ner, Oci, BM and Terp (up to 100 μM) on cell viability in PR and sensitive breast cancer cell lines. Figure 1 shows the concentration dependent effect of these terpenes in PR and sensitive MDA-MB-231 cells. The greatest effects on cell viability can be seen following treatment with Ner or BC (Figure 1A). BC had a relative IC_50_ of 27.6 μM and 4.4 μM in sensitive and resistant cell lines, respectively. Ner (Figure 1B) displays an IC_50_ of 26.3 μM and 27.1 μM in sensitive and resistant cell lines, respectively. In PR and sensitive MCF-7 cells a similar concentration-dependent effect can be seen (Supplemental Figure 1). BC (Supplemental Figure 1A) exhibits a relative IC_50_ of 2.5 μM and 10.8 μM, while Nerolidol (Supplemental Figure 1B) has an IC_50_ of 12.7 μM and 10.6 μM, in sensitive and resistant cells, respectively. Other terpenes screened showed less potent effects in all cell lines. In the MCF-10A non-tumorigenic breast epithelial cell line, no significant decrease in cell viability was observed for any of the five terpenes tested at concentrations of 10 or 100 μM (Supplemental Figure 2), suggesting the cytotoxic effects observed are specific to the cancer cell lines.

**FIG. 1.**
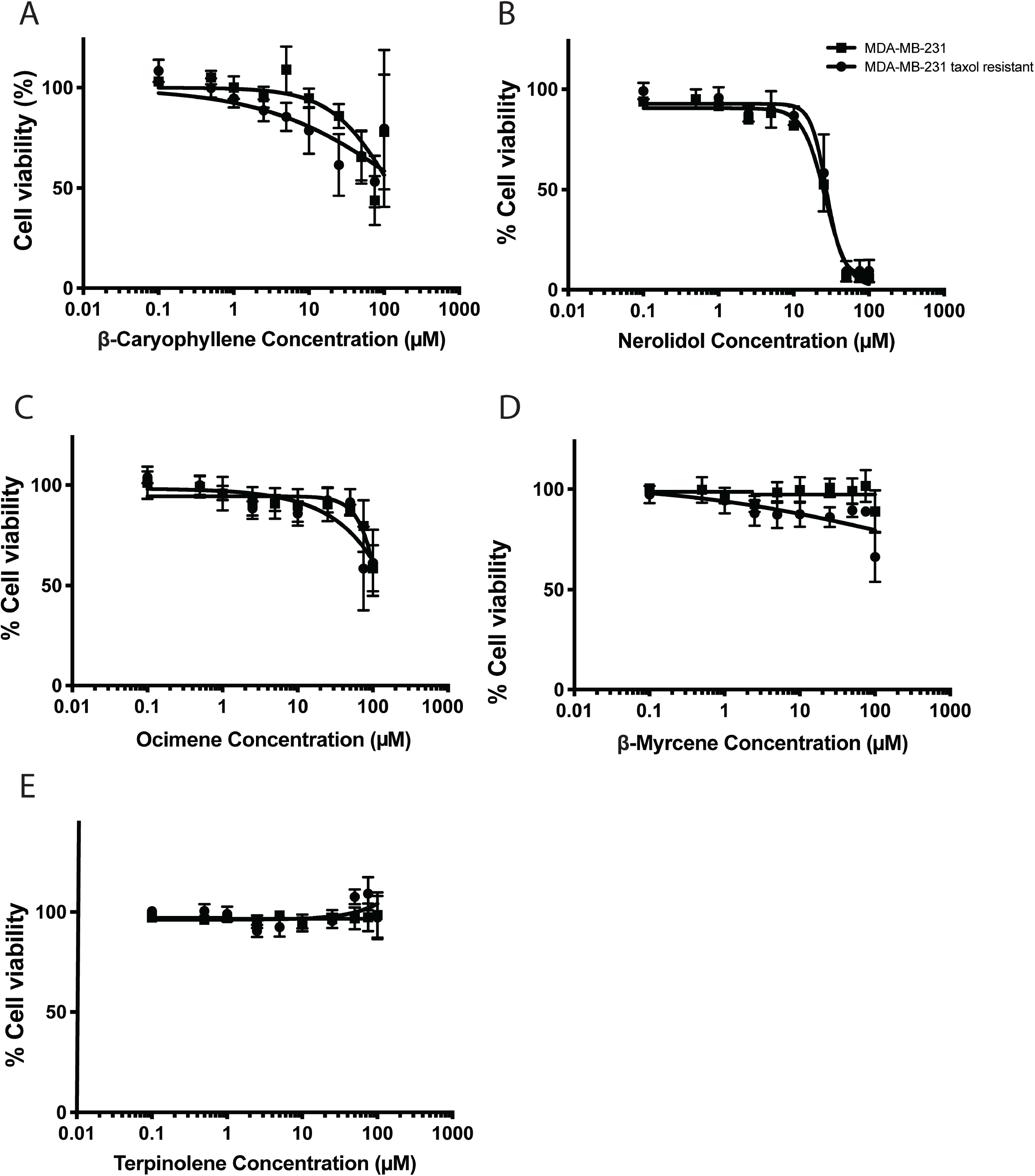
Effects of terpenes on cancer cell viability. Cell viability assays in taxol-resistant and sensitive MDA-MB-231 cells. Cells were treated with A: β-caryophyllene, B: Nerolidol C: Ocimene, D: β-myrcene E: Terpinolene at concentrations ranging from 0-100 μM. Cells were treated for 48h and fluorescence was detected after the addition of AlamarBlue reagent. n is at least 3 independent trials, represented as mean ± SEM.

### Apoptosis Detection

We next wanted to determine how the terpenes having the greater effects −Ner and BC–were decreasing cell viability. Since 5 and 10 μM had minimal impact on cell viability, these concentrations were chosen as to not affect the signal readouts and induce a bias. To explore if the cytotoxic effects of Ner or BC were related to the induction of apoptosis, an Annexin V assay was performed (Figure 2 A-B). MDA-MB-231 cells treated with 5μM Nerolidol showed a significant increase (p<0.0001) in Annexin V labelled cells with a mean of 48.5 % of cells when compared to vehicle control (mean of 10.2 %) (Figure 2A). MDA-MB-231 cells treated with 5 μM BC showed a significant increase (p<0.0001) in Annexin V labelled cells with a mean of 49.8 % of cells when compared to vehicle control (Figure 2A). There was no significant increase in propidium iodide labelled MDA-MB-231 cells after treatment with 5 μM Ner or BC (Figure 2A). To examine the apoptotic pathway further, a caspase 3/7 detection kit was used to evaluate caspase involvement. There was a significant increase in caspase3/7 detection in MDA-MB-231 cells treated with 10 μM Ner or BC (p<0.05) (Figure 2B). To further confirm the involvement of Caspase-3 in the apoptotic pathway induced by the terpenes, western blots of MDA-MB-231 cells were prepared following treatment with 5 μM Ner or BC (Figure 2C). All samples were probed for β–Actin as a loading control (Figure 2C). Caspase-3 can be seen at 32 kDa and its expression is lower in cells treated with Ner and BC, while the expression of cleaved caspase-3 at 11 kDa is increased (Figure 2C) following drug treatment.

**FIG. 2.**
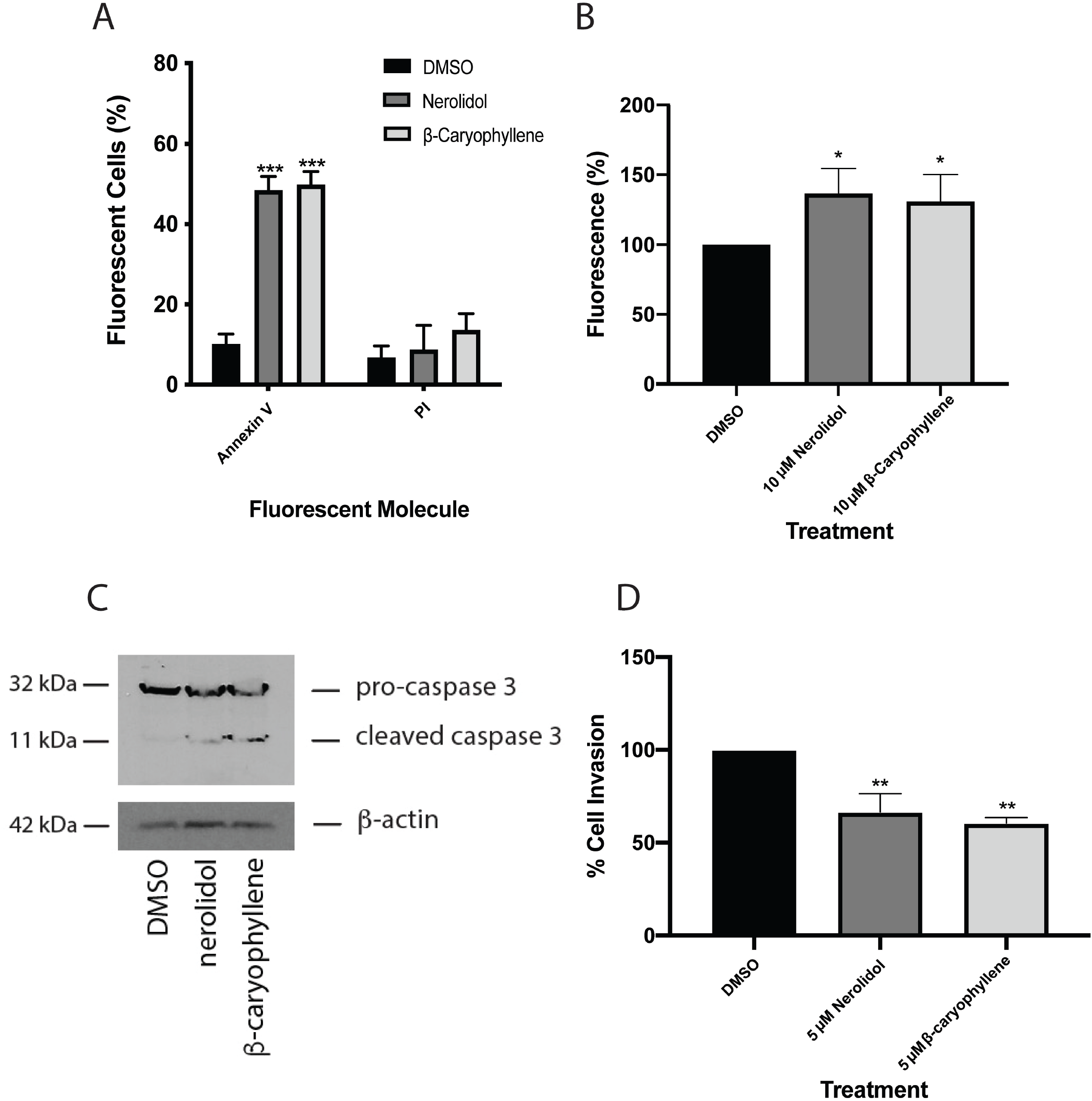
Effects of terpenes on apoptosis and cellular invasion. A: Histogram showing the % of annexin V-labeled MDA-MB-231 cells and % cells stained for propidium iodide. Cells were treated for 24 h with either the DMSO vehicle, Ner or BC B: Caspase 3/7 detection measured as percent fluorescence relative to DMSO control treated cells in MDA-MB-231 cells Cells were treated with 5 μM Ner, BC or DMSO as a control for 30 minutes. C: Western blot of MDA-MB-231 cells following 24 h treatment with DMSO, Ner, or BC was performed using an anti-caspase 3 antibody, and β-tubulin was included as a loading control. Figure is a representative blot of n = 3 experiments. D: MDA-MB-231 cells invaded through Matrigel Invasion Chambers. Cells were treated for 24 hours with 5 μM Ner, BC, or DMSO as a control. Cells on individual wells were counted and an average of two wells were calculated.

### Anti-Invasive Properties

To assess the anti-invasive properties of Ner and BC, a matrigel invasion assay was conducted (Figure 2D). There was a significant decrease (p<0.01) in cellular invasion of PR MDA-MB-231 cells after 24h treatment with 5 μM Ner with a mean of 66.6 % invasion compared to the vehicle control (Figure 2D). There was also a significant decrease (p<0.01) in cellular invasion of PR MDA-MB-231 cells after 24h treatment with 5 μM BC with a mean of 60.6 % invasion compared to the vehicle control (Figure 2D).

### Multidrug Resistance Mechanism Involvement

Using PR cell lines, we next wanted to see if the manipulation of efflux pumps could further enhance the cytotoxic effects of terpenes and provide insight into the mechanism involved in their cytotoxic activity. Verapamil can inhibit p-glycoprotein pump (p-gp) activity and was used in combination with terpene treatment to determine if it could enhance their effects. Verapamil alone did not have any significant effect on the cells used for this experiment. In PR MDA-MB-231 cells there was a significant decrease in cell viability between cells treated with Ner, BC, Oci, or BM plus verapamil when compared to the vehicle control and terpene alone, however these results were not significantly different from verapamil treatment alone (Figure 3A-E). Similar results were observed in PR MCF-7 cells (Supplemental Figure 3A-E). Because the combination of terpene and verapamil is not significantly different from both terpenes alone and verapamil alone, the combination is not providing enhanced effects and suggests the potential involvement of other efflux pumps or mechanisms of resistance that are not blocked by verapamil.

**FIG. 3.**
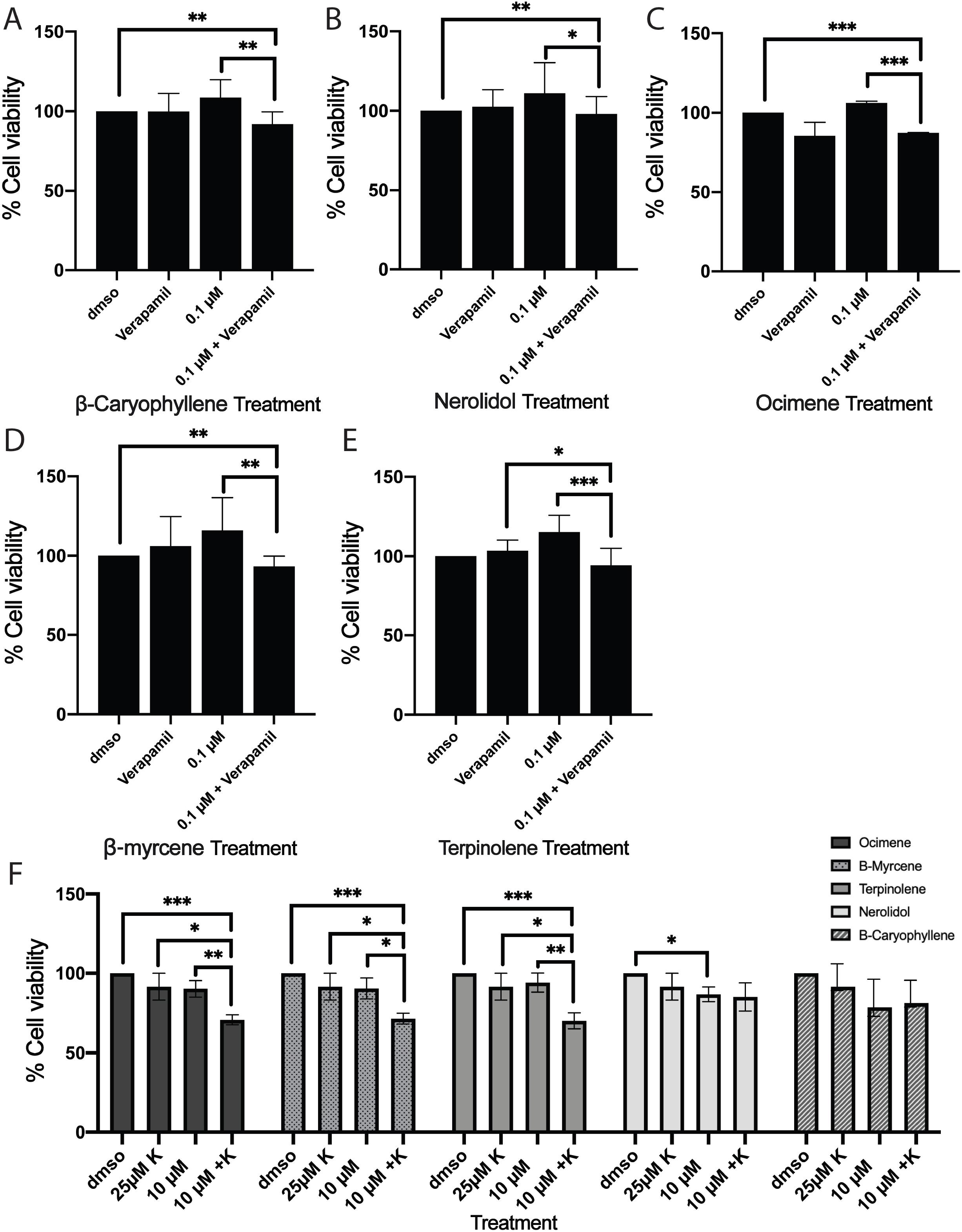
Combinations with efflux pump inhibitors. Verapamil’s effects on cell viability in MDA-MB-231 cells (A-E). Cells were treated with 0.1 μM DMSO, BC, Ner, Oci, BM, or Terp with or without 10 μM verapamil for 48h. Cell viability of taxol-resistant MDA-MB-231(F) cells after 48h treatment with 0.1-10 μM of Ner, BC, Oci, BM, or Terp in combination with 25 μM Kaempferol.

### Combination of Terpenes with Kaempferol

Flavonoids such as kaempferol, a compound found in cannabis, display properties related to the inhibition of efflux pumps such as MRP1. We measured the effect of kaempferol and terpenes, when used concomitantly, to identify whether kaempferol would affect cytotoxicity. The viability of PR MDA-MB-231 cells (Figure 3F) and PR MCF-7 cells (Supplemental Figure 3F) in the presence of the five terpenes alone or in combination with 25 μM kaempferol for 48 h was determined using experimental procedures like the ones described for the verapamil experiment. In both cell lines, kaempferol did not produce any significant effects on cell viability on its own when compared to the vehicle control (Figure 3F and Supplemental Figure 3F). In PR MDA-MB-231 cells, there was a significant difference in viability between 10μM Oci, BM, and Terp treatment in combination with 25μM kaempferol and vehicle control treatment, kaempferol alone, and terpene alone (Figure 3F). In PR MCF-7 cells there was a significant decrease in cell viability between 10 μM Oci and 25μM kaempferol combination treatment and vehicle control treatment, kaempferol alone, and terpene alone (Supplemental Figure 3F). These results suggest a potential effect of efflux pumps blocked by kaempferol in the cytotoxicity effects observed with Oci, BM, and Terp in PR MDA-MB-231 cells and Oci in PR MCF-7 cells.

### Synergy with Cannabinoids

As previous studies have shown cannabis have anti-cancer properties (Tomko et al., 2020), we wanted to evaluate if there could be greater cytotoxic effects observed when using the terpene showing the greatest effects, Ner, combined with the well-characterized cannabinoids THC or CBD. Following 48 h treatment with THC or CBD and Ner, the potential synergy, additivity, and antagonism of these compounds were measured in PR MDA-MB-231 and PR MCF-7 cell lines (Figure 4 and Supplemental Figure 4). Table 1 and Supplemental Table 1 show the maximum and minimum 3 concentration combinations that induced the highest or lowest levels of interaction for CBD and THC, respectively. A score <-10 is likely antagonistic (yellow); between −10 and +10 is likely additive (light yellow and light blue); >+10 is likely synergistic (blue). Our results show antagonism, additivity, and synergy can be seen in the combinations at different concentration combinations. In PR MDA-MB-231 cells our results indicate some synergy between Ner and THC (synergy score of 13) and more synergy between Ner and CBD (synergy score of 23) is occurring (Table 1). Much higher synergy scores were seen in PR MCF-7 cells where the combination of Ner and THC produced synergy scores up to 44 and Ner and CBD produced scores up to 27 (Supplemental Table 1). These results suggest that variable synergistic effects can be observed when Ner is combined with cannabinoids like THC or CBD and the magnitude of these results are cell-line dependent.

**FIG. 4.**
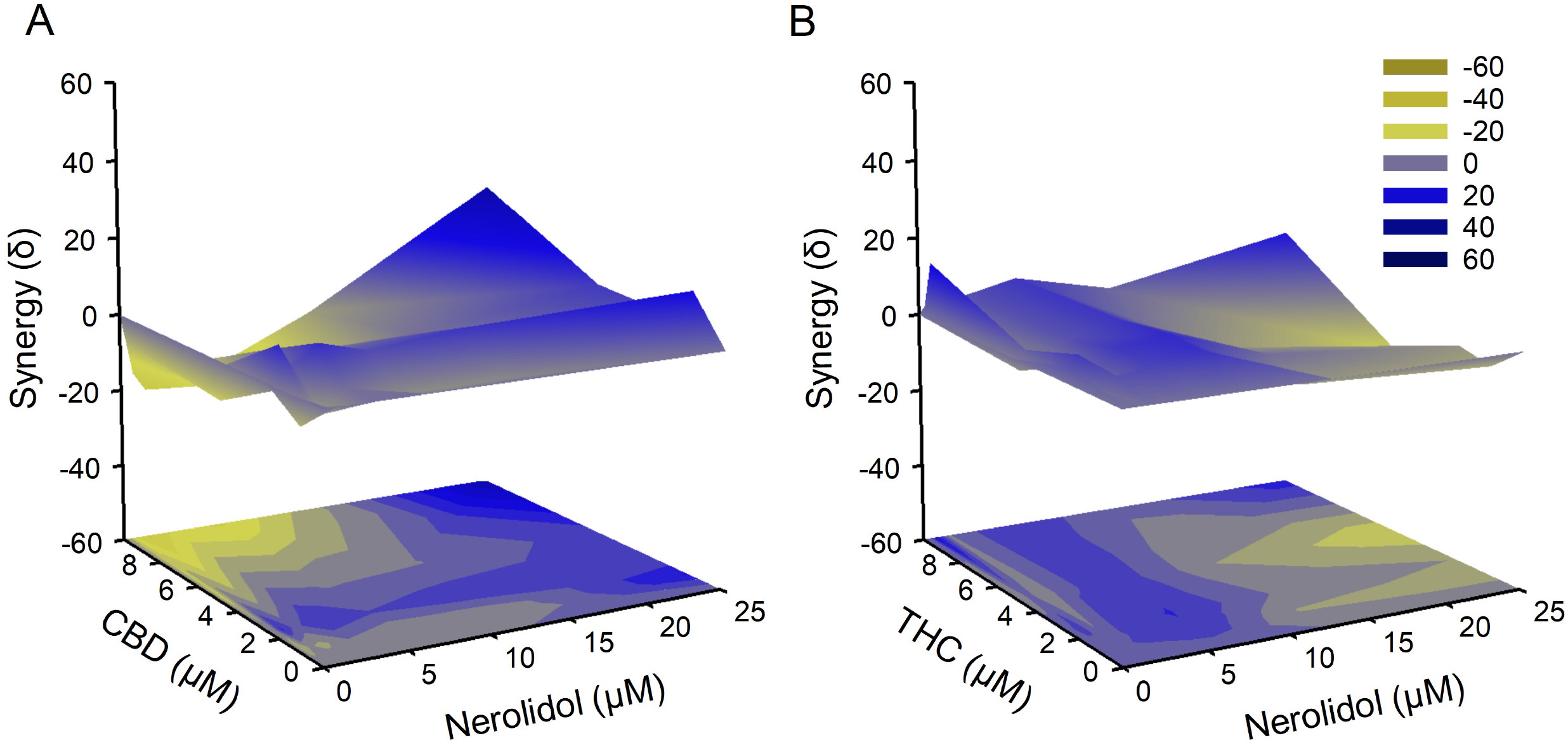
Assessment of synergy between Δ9-tetrahydrocannabinol or cannabidiol and Nerolidol. 2D and 3D synergy landscapes for the combinations of Δ9-tetrahydrocannabinol (A-B) or cannabidiol (C-D) (0-10μM) with Nerolidol (0-25μM) in taxol-resistant MDA-MB-231 cells. Yellow indicates areas of antagonism (synergy scores <-10), Blue indicates areas of synergy (synergy score >10), scores >-10 and <10 indicates areas of additivity.

**TABLE 1.**
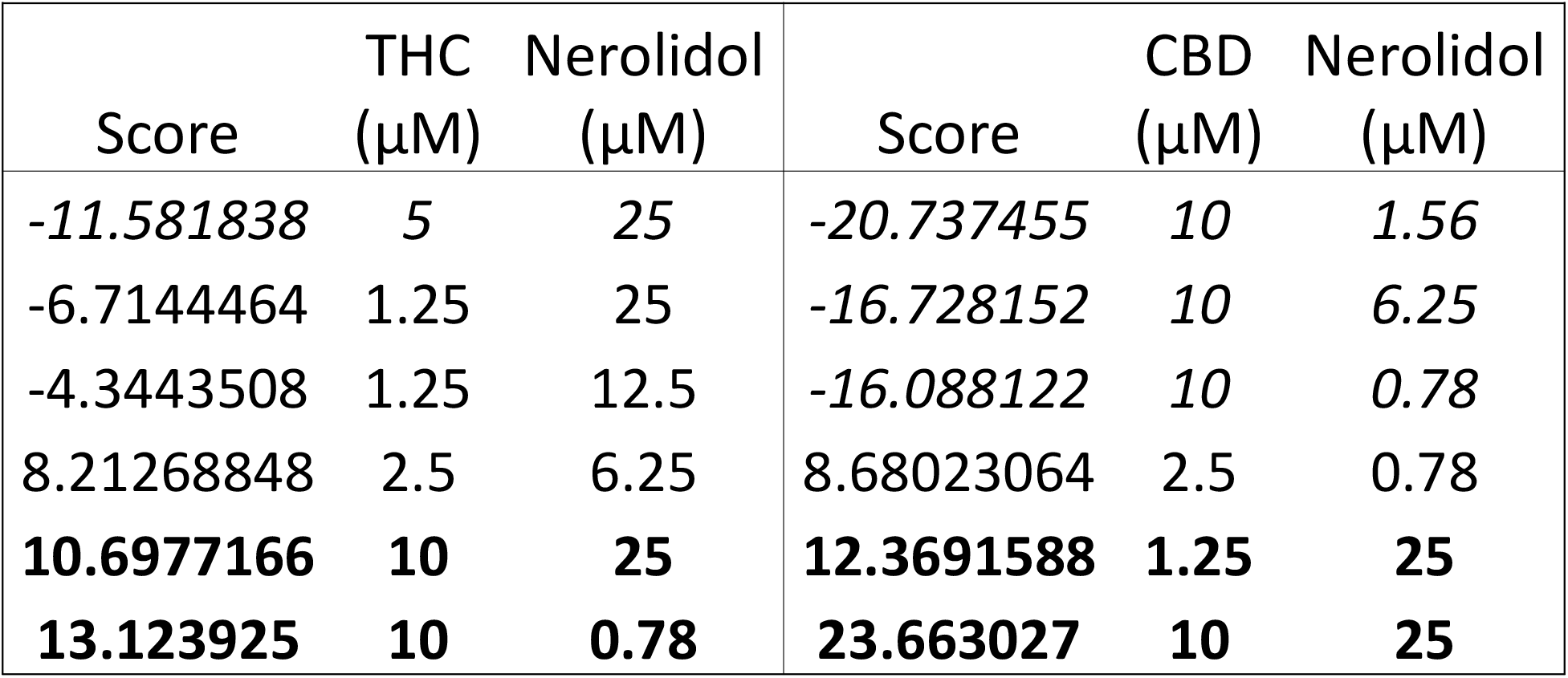
Highest and lowest levels of interaction between CBD, THC, and Nerolidol in taxol-resistant MDA-MB-231 cells. Italic values indicate antagonism (synergy score<10), bolded values indicate synergy (synergy score>10) white indicates additivity (synergy score >-10 and <10).

## Discussion

The effects of several terpenes found in cannabis, including Ner and BC, were evaluated for their anti-cancer potential in taxol-resistant breast cancer cell lines. Our results show that Ner and BC can reduce cell viability in taxol-resistant breast cancer cell lines, and that caspase-3 mediated apoptosis is involved in this reduction. We also demonstrate the Ner and BC are able to reduce the invasiveness of these cells. We further showed that anti-cancer effects were not enhanced when terpenes were combined with a known p-gp inhibitor, verapamil, but that some terpene effects were enhanced when combined with kaempferol, a flavonoid suggested to inhibit other efflux pumps such as MRP1 (Lee & Jeong, 2021). Finally, we tested for potential synergy between Ner and the well characterized cannabinoids THC and CBD and found that the highest levels of synergy are seen when Ner is combined with CBD.

Our results indicate that the terpenes tested produced a range of anti-proliferative effects in taxol-sensitive and resistant cell lines, with differences in their IC_50_. For example, while Ner and BC were both able to reduce cell viability to below 50%, Terp and BM were unable to achieve this reduction and had minimal effects at concentrations up to 100 μM. Oci was able to reduce cell viability to approximately 50%, however minimal effects were seen at lower concentrations. In contrast, Ner and BC both showed gradual concentration dependent decreases. This led us to focus our investigation on Ner and BC, finding that they both exhibited IC_50_ values ranging between 4.5 and 28 μM in sensitive and PR MDA-MB-231 cells and between 2.5 and 13 μM in sensitive and PR MCF-7 cells −lower than those previously reported in the literature (Hanušová et al., 2017). Due to the lower terpene content by weight present in cannabis (Chan et al., 2016) these concentrations may not be reached in breast tissue by consuming cannabis, however, various methods including targeted nanoparticles could be used to allow effective concentrations to be achieved in the breast tissue as suggested with other drugs (Tomko et al., 2020). Studies have shown that some terpenes including Ner and BC may exhibit their anti-cancer properties by inducing apoptosis through various pathways (Arul et al., 2020; Hanušová et al., 2017; Tomko et al., 2020). We show that in PR MDA-MB-231 cells, apoptosis is induced by Ner and BC and is mediated through a caspase-3 dependent pathway. In addition to apoptosis, other signaling pathways linked to cellular invasion are inhibited by Ner and BC, as a reduction in invasion of PR MDA-MB-231 cells was achieved by both compounds, suggesting these compounds could be used to help reduce invasiveness of breast cancer.

Previous research has suggested the potential of cooperative effects occurring between compounds found in cannabis and studies have tried to determine how different compounds in the plant may work together to exert their effects. One study in particular found that a botanical cannabis preparation showed greater anti-cancer effects in *in vitro* and *in vivo* breast cancer models compared to pure THC alone; the compounds present in the botanical preparation were not completely identified (Blasco-Benito et al., 2018). At the moment, little is known about which potential combinations could be used therapeutically and may produce synergistic effects when combined. Our approach investigated several compounds found in cannabis to identify if they could increase the effects shown by the cannabis terpenes chosen and how these were acting. Knowing that taxol-resistant cells possessed drug-resistant mechanisms implicating higher efflux pumps activity, we first evaluated if inhibiting the p-gp could increase the anticancer effects previously shown. After inhibiting the p-gp with a known inhibitor, we did not see further increases in cytotoxicity with the terpene treatments tested. We then evaluated if the cannabis flavonoid kaempferol, an inhibitor for other efflux pumps such as MRP1, could increase the cytotoxic effects of the terpenes (Lee & Jeong, 2021) and found that kaempferol cotreatment with the terpenes Oci, BM, and Terp significantly decreased cell viability. The lack of effect of verapamil and the effects of kaempferol suggest that MRP1, rather than the p-gp, may be involved in the efflux of several terpenes in these PR cells. Another explanation for kaempferol’s effects could be through its interactions with Glutathione S-transferases (GST). GSTs play a role in detoxifying xenobiotics and allowing their export from the cell through GS-X pumps, often overexpressed in resistant cancers. Kaempferol has been shown to inhibit GSTs, specifically M1-1 and M2-2, which could result in a reduction of drug efflux and improved anticancer effects (Hayeshi et al., 2007).

Our assessment of potential synergy between Ner and THC or CBD found that low levels of synergy are present between Ner and both cannabinoids and some higher levels of synergy are seen between Ner and CBD. Although studies to date have yet to demonstrate synergy between cannabinoids and terpenes using isolated compounds, combinations of compounds from cannabis and anti-cancer agents have shown improved effects compared to the compounds alone (Cao et al., 2019; Fraguas-Sánchez et al., 2020; Hanušová et al., 2017; Legault & Pichette, 2007). Our results suggest that flavonoids such as kaempferol could further increase the levels of synergy if combined with other compounds, due to its potential effect on reported drug-resistance mechanisms.

Our findings indicate that there is the potential for flavonoids, cannabinoids, and terpenes to act together to produce greater anti-cancer effects. These results have yet to be validated *in vivo*, however they suggest that terpenes found in cannabis should be further studied to identify their potential inclusion in treatment of drug-resistant breast cancer. More investigation is needed to determine the exact mechanisms of action of how these terpenes may act in concert with flavonoids, cannabinoids and other chemotherapeutic agents to treat chemotherapy-resistant breast cancer.

