## supplemental material for "Anti-cancer potential of cannabis terpenes in a taxol-resistant model of breast cancer"

Tomko, A. Supplemental Table 1

**SUP TABLE 1. Highest and lowest levels of interaction between CBD, THC, and Nerolidol in taxol-resistant MCF-7 cells.**
Italic values indicate antagonism (synergy score<10), bolded values indicate synergy (synergy score>10) white indicates additivity (synergy score >-10 and <10).

| Score | THC (µM) | Nerolidol (µM) | Score | CBD (µM) | Nerolidol (µM) |
|---|---|---|---|---|---|
| *-42.745557* | *5* | *25* | *-47.879118* | *1.25* | *25* |
| *-41.196531* | *2.5* | *25* | *-32.789706* | *1.25* | *3.12* |
| *-40.342722* | *1.25* | *25* | *-23.676037* | *5* | *25* |
| **13.9353743** | **5** | **12.5** | **23.5151208** | **2.5** | **25** |
| **37.2572171** | **10** | **25** | **27.1080452** | **5** | **1.56** |
| **44.8817469** | **10** | **12.5** | **27.2959726** | **5** | **6.25** |

Tomko, A. Supplemental Figure 1

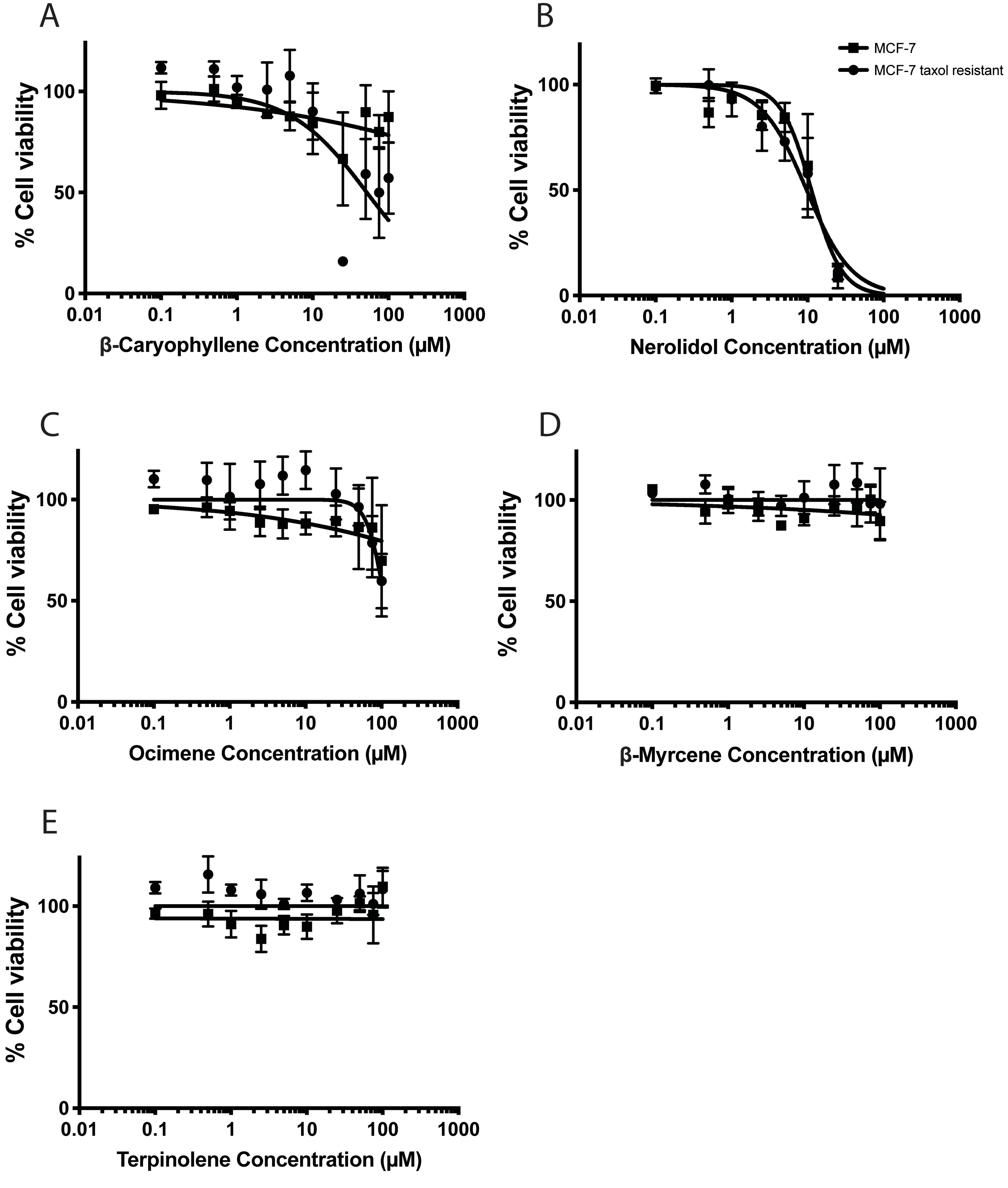


**SUP FIG. 1. Effects of terpenes on cancer cell viability**

Cell viability assays in taxol-resistant and sensitive MCF-7 cells. Cells were treated with A: β-caryophyllene, B: Nerolidol C: Ocimene, D: β-myrcene E: Terpinolene at concentrations ranging from 0-100 μM. Cells were treated for 48h and fluorescence after the addition of AlamarBlue reagent. N is at least 3 independent trials, represented as mean ± SEM.

Tomko, A. Supplemental Figure 2

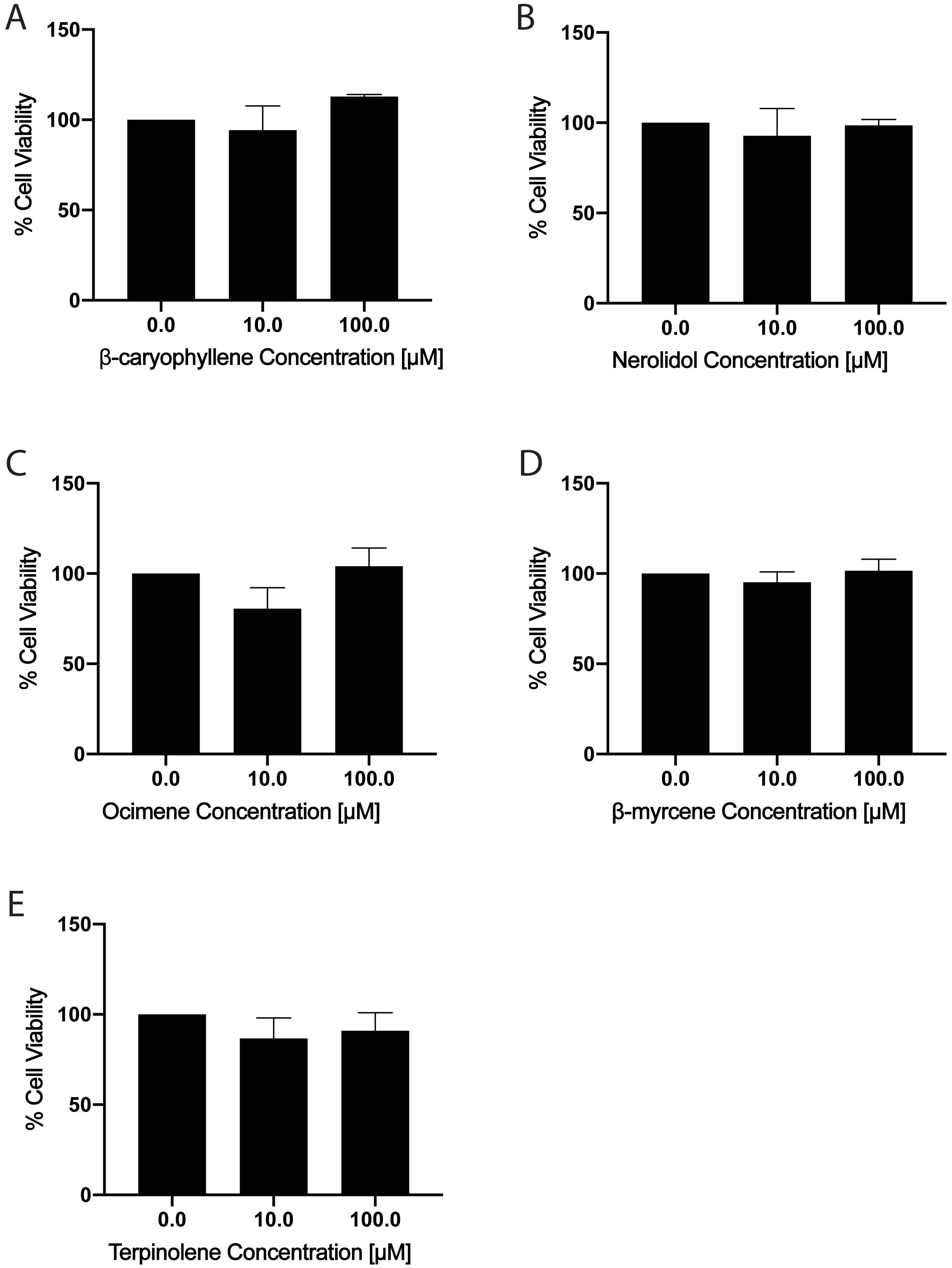


**SUP FIG. 2. Effects of terpenes on non-tumorigenic cell viability**

Cell viability assays in MCF10-A cells. Cells were treated with A: β-caryophyllene, B: Nerolidol C: Ocimene, D: β-myrcene E: Terpinolene at concentrations ranging from 0-100 μM. Cells were treated for 48h and fluorescence was detected after the addition of AlamarBlue reagent. N is at least 3 independent trials, represented as mean ± SEM.

Tomko, A. Supplemental Figure 3

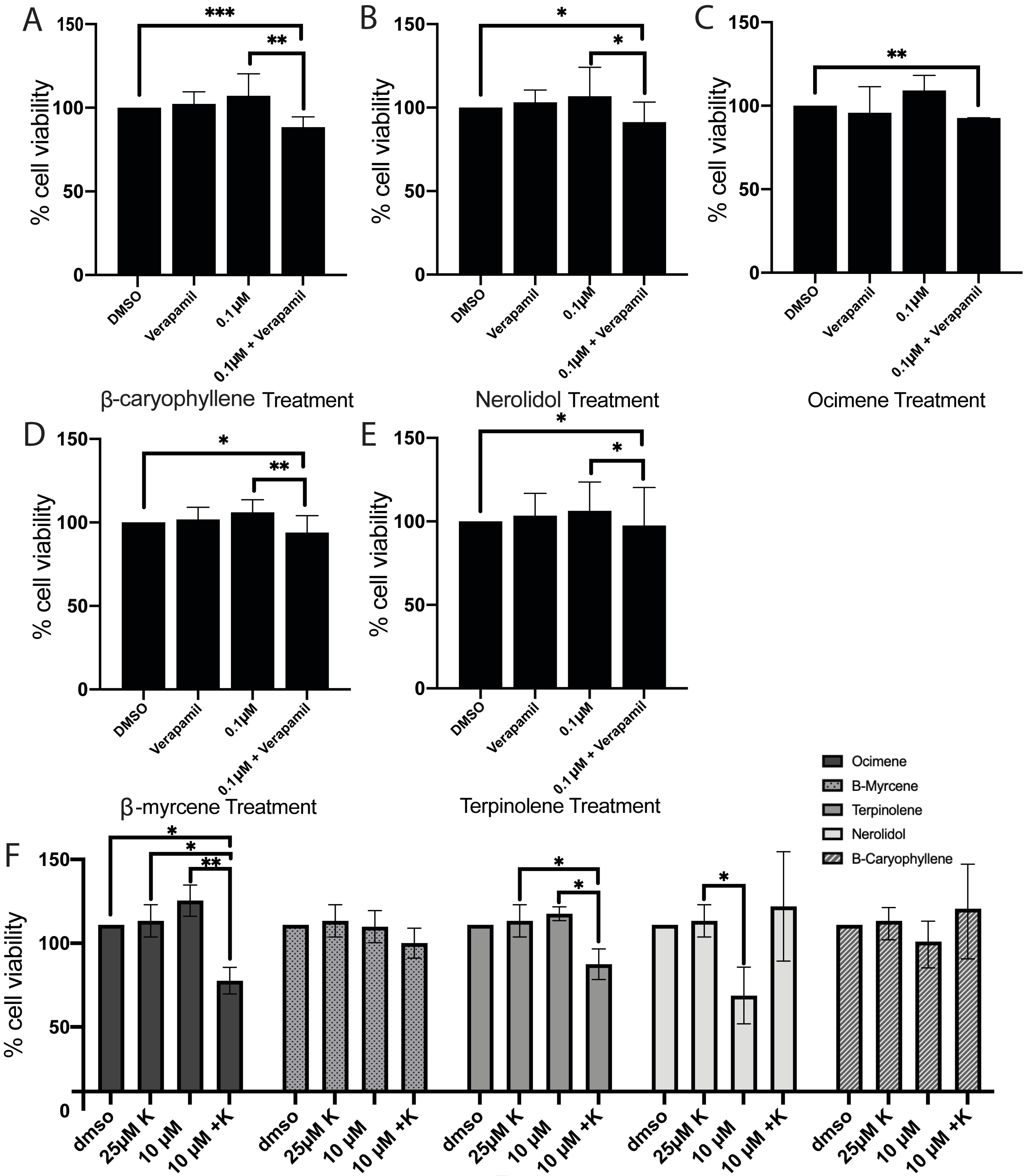


**SUP FIG. 3. Combinations with efflux pump inhibitors**

Verapamil induced a decrease in cell viability in MCF-7 (A-E) taxol-resistant cells. Cells were treated with 0.1 µM DMSO, BC, Ner, Oci, BM, or Terp with or without 10µM verapamil for 48h. Cell viability of taxol-resistant MCF-7 (F) cells after 48h treatment with 0.1-10 µM of Ner, BC, Oci, BM, or Terp in combination with 25 µM Kaempferol.

Tomko, A. Supplemental Figure 4

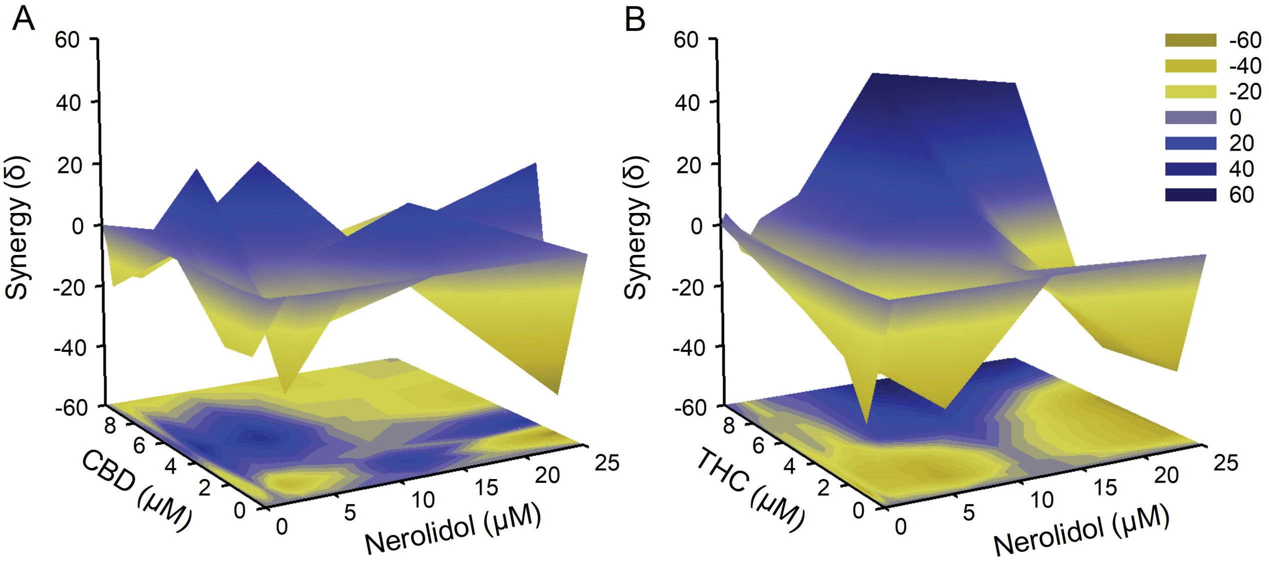


**SUP FIG. 4. Assessment of synergy between Δ9-tetrahydrocannabinol or cannabidiol and Nerolidol**
Assessment of synergy between Δ9-tetrahydrocannabinol or cannabidiol and Nerolidol. 2D and 3D synergy landscapes for the combinations of Δ9-tetrahydrocannabinol (A-B) or cannabidiol (C-D) (0-10 μM) with Nerolidol (0-25 μM) in taxol-resistant MCF-7 cells. Yellow indicates areas of antagonism (synergy score<-10), Blue indicates areas of synergy (synergy score>10), scores >-10 and <10 indicates areas of additivity.
